# Distinct neural representations during a brain-machine interface and manual reaching task in motor cortex, prefrontal cortex, and striatum

**DOI:** 10.1101/2023.05.31.542532

**Authors:** Ellen L. Zippi, Gabrielle F. Shvartsman, Nuria Vendrell-Llopis, Joni D. Wallis, Jose M. Carmena

## Abstract

Although brain-machine interfaces (BMIs) are directly controlled by the modulation of a select local population of neurons, distributed networks consisting of cortical and subcortical areas have been implicated in learning and maintaining control. Previous work in rodent BMI has demonstrated the involvement of the striatum in BMI learning. However, the prefrontal cortex has been largely ignored when studying motor BMI control despite its role in action planning, action selection, and learning abstract tasks. Here, we compare local field potentials simultaneously recorded from the primary motor cortex (M1), dorsolateral prefrontal cortex (DLPFC), and the caudate nucleus of the striatum (Cd) while nonhuman primates perform a two-dimensional, self-initiated, center-out task under BMI control and manual control. Our results demonstrate the presence of distinct neural representations for BMI and manual control in M1, DLPFC, and Cd. We find that neural activity from DLPFC and M1 best distinguish between control types at the go cue and target acquisition, respectively. We also found effective connectivity from DLPFC→M1 throughout trials across both control types and Cd→M1 during BMI control. These results suggest distributed network activity between M1, DLPFC, and Cd during BMI control that is similar yet distinct from manual control.

## Introduction

The ability to volitionally modulate the activity of neurons has been demonstrated in both human subjects and animals [1]. This skill is fundamental to the operation of brain-machine interfaces (BMIs), in which external devices, such as computer cursors or robotic arms, are controlled by the production of specific patterns of neural activity. Motor BMIs specifically rely on the modulation of motor cortical neural activity to control external devices [2]–[4], while also relying on higher-level cognitive processes to integrate sensory information, plan and initiate actions, and monitor visual feedback of the effector [5]. These systems hold great potential for rehabilitation and restoration of lost motor function [4], [6], but the underlying neural processes involved in controlling a motor BMI are still unclear.

Controlling a BMI shares characteristics with both overt motor control and abstract cognitive tasks [7]–[9]. BMI control requires users to produce specific patterns of neural activity to manipulate their physical environment without physical movement and often without proprioceptive feedback. Thus, BMI control may rely on networks underlying both overt motor control and abstract cognitive tasks. While both BMI control and overt motor control rely on the direct modulation of the motor cortex, the mapping between neural activity and output is distinct between these types of control because a specific subpopulation of neurons is selected for BMI control. The prefrontal cortex and dorsal striatum are important for both motor control [10]–[13] and for learning abstract associations [14]–[17].

The primary motor cortex (M1), the dorsolateral prefrontal cortex (DLPFC) and the caudate nucleus of the striatum (Cd) are extensively interconnected with one another, as well as with other sensory and higher-level associational areas involved in goal-directed behavior [18]–[22]. Cortico-striatal plasticity is necessary for motor control [23] and cortico-striatal and cortico-cortical interactions are involved in abstract skill learning [18], [24], [25], suggesting that these interactions may play an important role in BMI control.

Successful control of a BMI depends on multiple cognitive processes that are employed differentially across events within the task. For example, in a center-out BMI task, the go cue may be more dependent on action selection and initiation, while control at target acquisition may depend more on error correction and fine movement control. Therefore, it is possible that different regions of interest (ROIs), such as motor cortex, prefrontal cortex, or striatum, are more or less involved at different points within a trial. For instance, the prefrontal cortex may be more involved at the go cue when action planning and selection are critical [13], [18], [26] whereas the primary motor cortex may be more involved at target acquisition when movement execution and fine effector control are more important [27]. Thus, it is important to investigate neural activity in each ROI at multiple task events to better understand its role during manual and BMI control.

Previous studies that have used BMIs to dissect cognitive processes have primarily focused on neural activity in BMI-selected neurons, motor cortical neurons whose activity is directly input to the BMI [28]. A few studies have investigated the role of neural activity from non-BMI neurons within the motor cortex [29], [30], highlighting that BMI control involves large-scale networks that extend beyond BMI-selected neurons. However, it is becoming apparent that extending these investigations to brain areas beyond the motor cortex will be essential for gaining a more complete understanding of BMI control [31]. Previous studies have shown the involvement of cortico-striatal networks during a BMI task in rodents [32]–[34] and of the prefrontal cortex in BMI skill acquisition in humans [8], [35], [36]. However, due to experimental constraints, these studies relied on simplified, one-dimensional BMI tasks, making it difficult to gain insight into the distributed network activity that may be specific to more complex and cognitively demanding two-dimensional (2-D) BMI control. To overcome these limitations, we successfully trained non-human primates (NHPs) to perform a 2-D BMI center-out task with recordings from a semi-chronic microdrive. These microdrives have previously been used to study large-scale networks [37], [38], but never in the context of BMI control. To our knowledge, this is the first time recordings from a semi-chronic microdrive have been used to drive a BMI in NHPs, allowing us to obtain simultaneous cortical and subcortical recordings to make direct comparisons of activity in these regions and investigate their interactions during BMI control.

Here, we use this unique experimental opportunity to investigate the modulation of and interactions between M1, DLPFC, and Cd activity during a motor BMI task and compare this activity to that during manual (overt motor) control. We perform these analyses at both the go cue and at target acquisition allowing us to compare the role of each ROI across two task events that involve distinct cognitive processes. We demonstrate that DLPFC holds the most information for distinguishing BMI and manual control at the go cue, while M1 is most informative at target acquisition. Additionally, we show that directed information flow is present from DLPFC→M1 in both BMI and manual control and from Cd→M1 during BMI control. These directed interactions were present at both the go cue and at target acquisition. Ultimately, our findings confirm the involvement of distributed cortical and subcortical networks during BMI control and identify similar but distinct network activity during manual control.

## Results

To explore whether there are BMI-specific neural representations in M1, DLPFC, and Cd, we simultaneously recorded from these regions during a BMI control task, a manual control task, and a baseline rest period. Two rhesus macaques (Monkey H and Monkey Y) were implanted with a custom-fit large-scale semi-chronic microdrive array on the left hemisphere (Figure 1a). Single- and multi-unit recordings from the motor cortex were used as input to a BMI decoder, while local-field potentials (LFP) were simultaneously recorded from three regions of interest (ROIs): M1, DLPFC, and Cd. Each day, the animals performed a two-dimensional, self-initiated, center-out task, in which they were instructed to move a cursor from a center target to one of eight pseudo-randomly instructed peripheral targets for a juice reward. A successful center-out trial required a brief hold at the center target, moving to the peripheral target within a specified time, and a brief hold at the target (Figure 1b). First, they performed the task under manual control (Figure 1c), followed by a four-minute baseline period (Figure 1d). Then, they performed the same center-out task under BMI control (Figure 1e-f).

**Figure 1.**
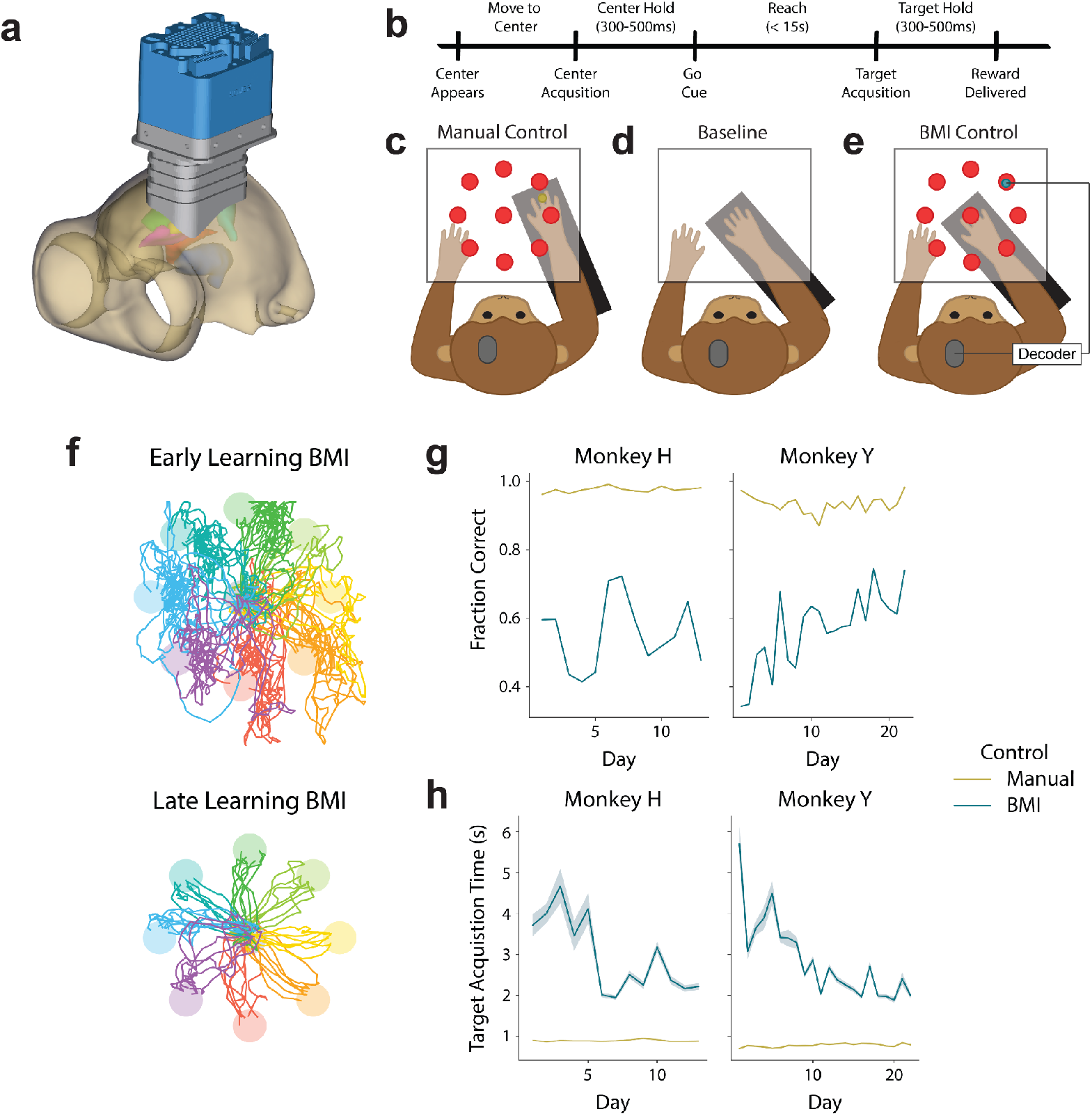
Experimental set-up and task performance. (a) A 3-D model of a custom-fit, large-scale, semi-chronic microdrive from Gray Matter Research used for simultaneous recordings of neural activity from ROIs at different depths. (b) Timeline of the center-out task (see Methods for details). (c) Two monkeys completed a 2-D, self-initiated, center-out movement task under manual control. (d) During a baseline period, neural activity was recorded during the absence of visual cues and the subject’s arm locked in a fixed position. (e) During BMI control, the subject’s arm was locked in a fixed position while they modulated neural activity to control a cursor in the same 2-D, self-initiated, center-out task. (f) Example BMI control trajectories from early learning (right) and late learning (left) to each target from Monkey Y. (g) Fraction of self-initiated trials that were successful in BMI control (teal) and manual control (yellow) for Monkey H (left) and Monkey Y (right). (h) Time from go cue to target acquisition in BMI control (yellow) and manual control (teal) for Monkey H (left) and Monkey Y (right). Shading represents the standard error of the mean across trials within a day.

### Successful BMI control with a semi-chronic array

Both monkeys successfully learned to perform the eight-target, center-out task using a cursor under BMI control. Task performance improved across days, despite using different direct units and decoders each day (see Methods for details). Although only one monkey significantly increased the fraction of successfully completed trials (Figure 1g; Linear regression, Monkey H: R^2^ = 0.003, p = 0.852, Monkey Y: R^2^ = 0.593, p = 2.78e-5), both monkeys learned to produce faster, straighter cursor trajectories, resulting in a decreased target acquisition time over days (Figure 1h; Linear regression, Monkey H: R^2^ = 0.035, p = 3.20e-27, Monkey Y: R^2^ = 0.093, p = 4.67e-141). This improved performance demonstrates that the animals were able to learn to control a BMI using neural activity recorded from a semi-chronic microdrive.

### Spectral power from M1, DLPFC, and Cd distinguishes BMI control, manual control, and baseline

To understand whether activity from M1, DLPFC, and Cd differs between BMI control, manual control, and baseline, we compared the spectral power in 5 distinct frequency bands across these three conditions (Supplementary Figures 1-2, see Methods for details). Using the LFP recorded from each ROI, we computed the mean power across electrode channels in each frequency band. For BMI and manual control, power was computed in the 500 ms window following the go cue and the 500 ms window preceding target acquisition on successful trials. The same calculations were performed in random non-overlapping 500ms windows during the baseline period. The most notable differences occurred between baseline and the two control tasks in the theta and beta frequency bands (Figure 2). At the go cue, theta power at baseline was significantly lower than during both manual and BMI control across all ROIs, while beta power at baseline was significantly higher than during either of the two control tasks (Figure 2a). At target acquisition, beta power at baseline remained significantly higher than during the two control tasks in M1 but was less consistent at other ROIs and frequency bands (Figure 2b). While these differences provide insight into how spectral power differs between the baseline rest period and the two control tasks at different task events, differences in spectral power within individual frequency bands between the two types of control were less consistent across animals.

**Figure 2.**
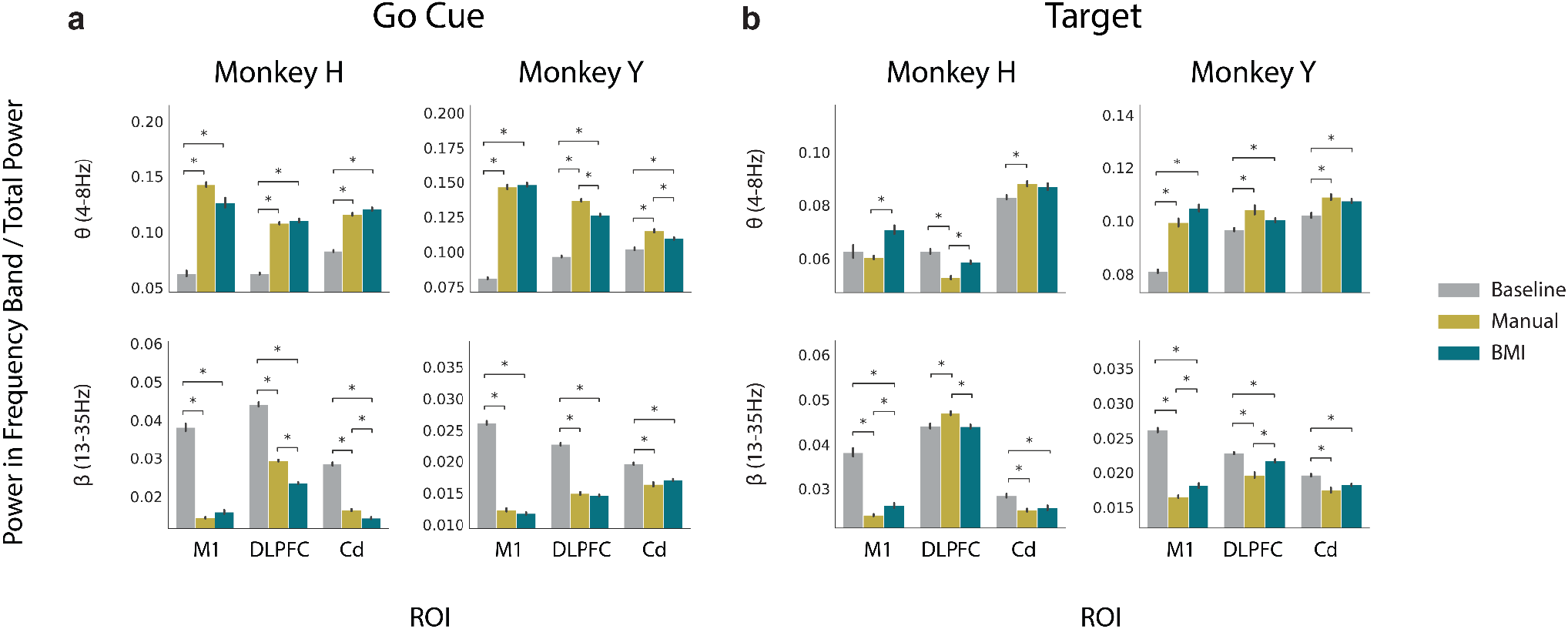
Theta and beta power in M1, DLPFC, and Cd is significantly different between baseline rest period and manual or BMI control. (a) Average theta (top) and beta (bottom) power normalized to total power in each ROI during baseline (gray), manual (yellow), and BMI (teal) at the go cue for Monkey H (left) and Monkey Y (right). Error bars represent standard error mean across days. (b) Same as (a), but for power features obtained at target acquisition rather than at the go cue. Normalized spectral power estimates significantly differing between task types after Bonferroni correction for multiple comparisons are indicated with an asterisk.

To determine whether combinations of spectral power from multiple frequency bands across our ROIs yield more distinct representations between task types, we trained linear discriminant analysis (LDA) classifiers to distinguish between these task types (Figure 3a). Although only theta and beta power showed consistent significant differences across task types, incorporating power from all frequency bands as input features yielded a higher classification accuracy than including any individual feature for each individual ROI, so all power features were included for each ROI in subsequent analyses (Supplementary Figure 3). We compared neural activity recorded during BMI control to activity recorded during both manual control and baseline, allowing us to identify task-related activity that is specific to BMI control (Figure 3a). At both the go cue and target acquisition, classifiers using power features from all ROIs, M1 + DLPFC + Cd, yielded the highest classification accuracy (Figure 3b-c, Supplementary Figure 4a-b). At target acquisition, classifiers that incorporated M1 power features (M1 + DLPFC, M1 + Cd, and M1 only) yielded significantly higher classification accuracy than classifiers without M1 power features (DLPFC + Cd, DLPFC only, and Cd only) (Supplementary Figure 4b). With BMI performance improving across days (Figure 1g-h), we considered whether these results varied across different stages of BMI learning. We repeated these analyses within early (first third of recording days; Monkey H: n = first 4 days, Monkey Y: n = first 7 days) and late learning periods (last third of recording days; Monkey H: n = last 4 days, Monkey Y: n = last 7 days). Both the go cue and target acquisition results were similar across BMI learning stages (Supplementary Figure 5). Overall, these results indicate that it is possible to distinguish between BMI control, manual control, and baseline using spectral power features from M1, DLPFC, and Cd. Furthermore, the increase in classification accuracy resulting from the inclusion of M1 activity indicates that there are distinct neural representations of the three task types in this region at target acquisition.

**Figure 3.**
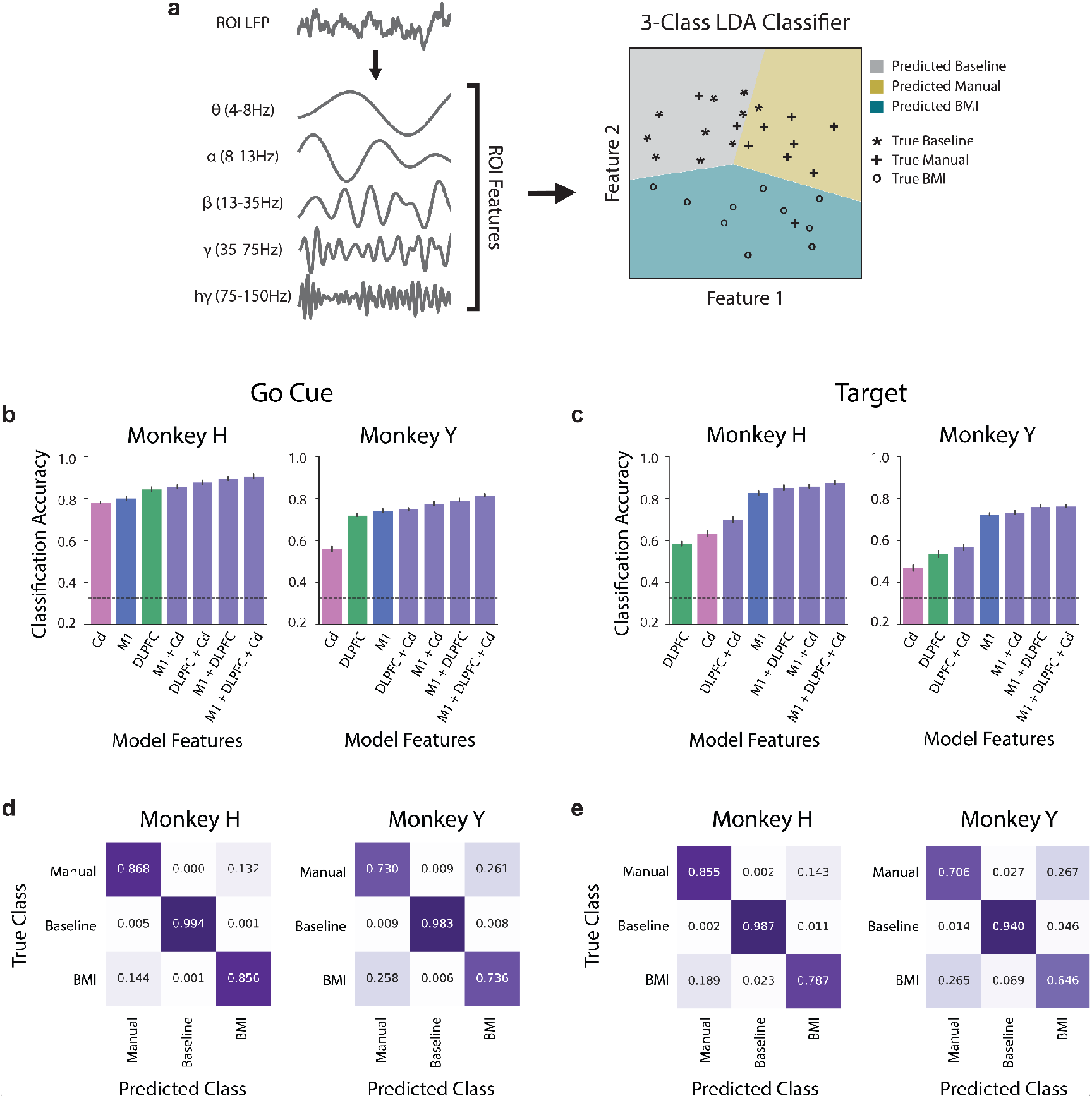
Spectral power in M1, DLPFC, and Cd distinguishes between BMI control, manual control, and baseline at the go cue and at target acquisition. (a) LFP was decomposed into 5 distinct frequency bands. The average power in each of these bands in each ROI was used as input to the classifiers. *3-class* LDA classifiers were trained to distinguish between BMI control, manual control, and baseline. (b) Mean 10-fold cross-validated classification accuracy across days for a *3-class* LDA classifier using all frequency bands from individual ROIs (M1 in blue, DLPFC in green, Cd in pink) or combinations of ROIs (purple) at the go cue for Monkey H (left) and Monkey Y (right). Classifiers are presented in order of ascending accuracy. Chance accuracy shown as a dashed line. Error bars represent standard error mean across days. (c) Same as (b), but for power features obtained at target acquisition rather than at the go cue. (d) Most misclassifications occurred between BMI control and manual control. Confusion matrix for the highest accuracy model at the go cue, which included all frequency bands from all ROIs. Numbers and shading correspond to the fraction of correct predictions within a class for Monkey H (left) and Monkey Y (right). (e) Same as (d), but for power features obtained at target acquisition rather than at the go cue.

**Figure 4.**
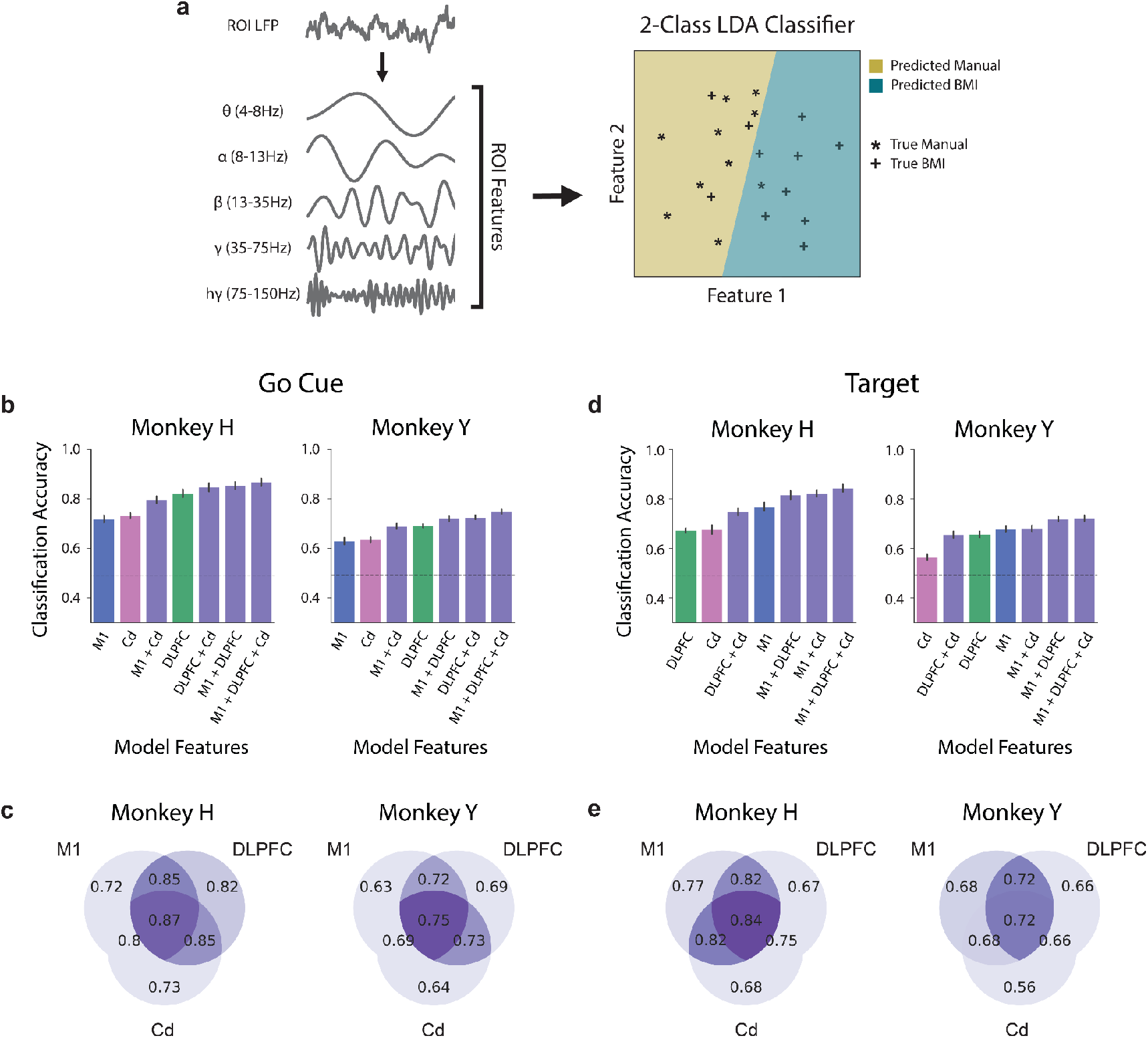
DLPFC activity best distinguishes between BMI and manual control at the go cue, while M1 activity best distinguishes between BMI and manual control at target acquisition. (a) LFP was decomposed into 5 distinct frequency bands. The average power in each of these bands in each ROI was used as input to the classifiers. *2-class* LDA classifiers were trained to distinguish between BMI control and manual control only. (b) Mean 10-fold cross-validated classification accuracy across days for a *2-class* LDA classifier trained to distinguish between BMI and manual control using all frequency bands from individual ROIs (M1 in blue, DLPFC in green, Cd in pink) or combinations of ROIs (purple) at the go cue for Monkey H (left) and Monkey Y (right). Classifiers are presented in order of ascending accuracy. Chance accuracy shown as a dashed line. Error bars represent standard error mean across days. (c) Mean classification accuracy for each combination of ROIs at the go cue. (d-e) Same as (b-c), but for power features obtained at target acquisition rather than at the go cue.

**Figure 5.**
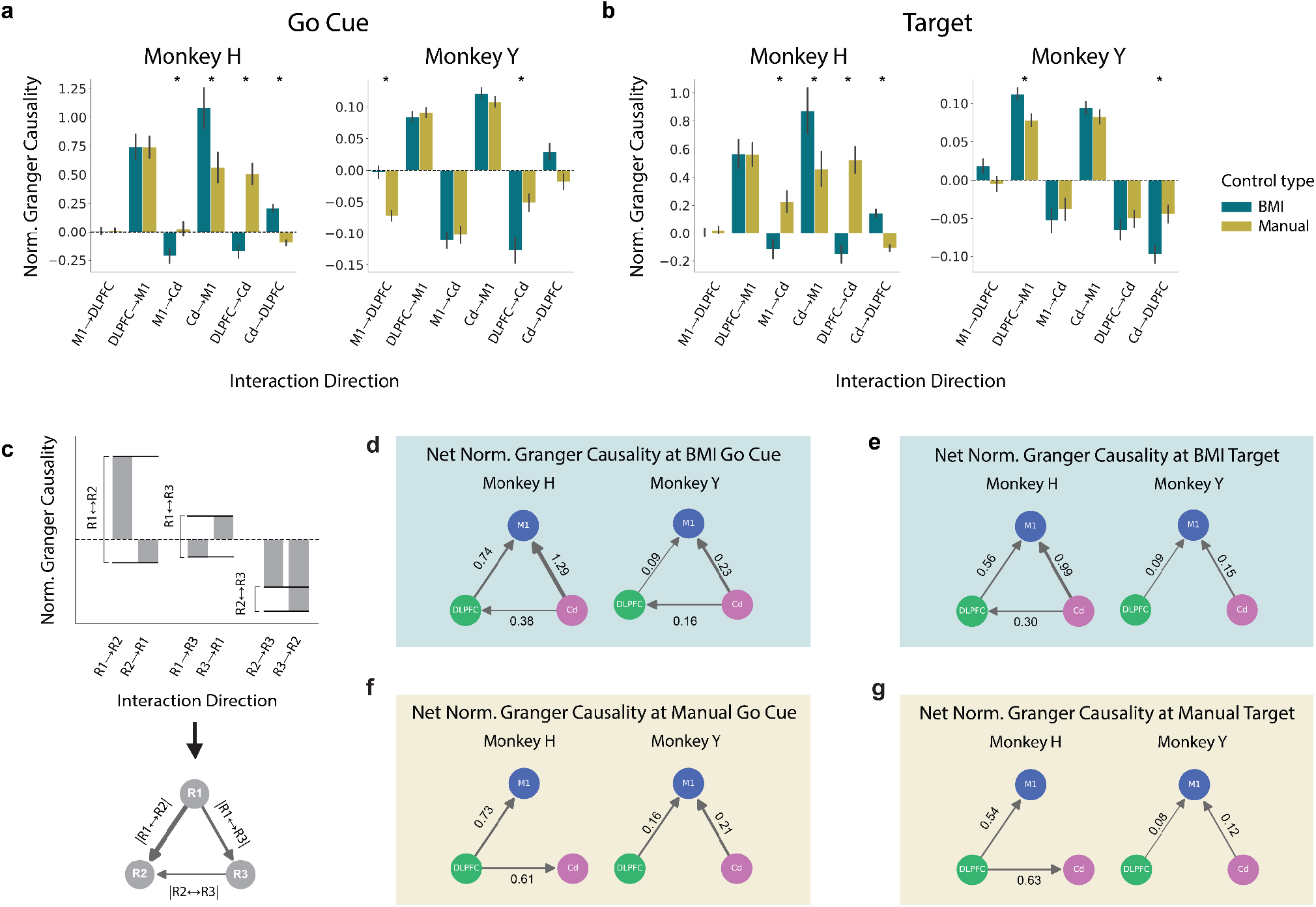
Net flow of information between M1, DLPFC, and Cd relative to baseline is similar across BMI and manual control. (a) Normalized Granger causality estimates at the go cue during BMI (teal) and manual (yellow) control. Normalized Granger causality greater than zero indicates an increase from baseline, whereas less than zero indicates a decrease from baseline. Normalized Granger causality estimates significantly differing between BMI and manual control are indicated with an asterisk (see Supplementary Table 4). Error bars represent standard error mean across days. (b) Same as (a) but for target acquisition, rather than go cue. (c) Schematic depicting the calculation of net normalized granger causality. The difference in normalized granger causality for reciprocal interactions was used to estimate net flow of information relative to baseline. (d) Significantly directed net normalized Granger causality at the go cue during BMI control (see Supplementary Table 5 for all values of net normalized Granger causality). (e) Same as (d), but for target acquisition during BMI control. (f) Net normalized granger causality at the go cue during manual control. (g) Same as (f), but for target acquisition during manual control.

To further explore the differences in the neural representations of BMI control, manual control, and baseline in M1, DLPFC, and Cd, we computed confusion matrices for the highest-accuracy classifiers, which were trained using power features from all ROIs (Figure 3d-e). Here, we assessed which type of misclassifications were most common. If the majority of misclassifications occur between BMI control and baseline, this would imply that these two classes are more similar to one another and distinct from manual control. Thus, we would infer that the differences in neural representations primarily correspond to the presence or absence of large, physical arm movements. On the other hand, a majority of misclassifications occurring between BMI control and manual control could indicate that differences in neural representations were largely task-related. Thus, quantifying the *3-class* LDA misclassifications allowed us to identify whether the model performed better at separating movement-related conditions or task-related conditions. At both the go cue and target acquisition, we found that the primary misclassifications occurred between BMI control and manual control, demonstrating that neural activity during the two tasks is more similar to one another than it is during baseline. This suggests that the primary distinction in neural representations was due to the presence or absence of a task, rather than the presence or absence of movement.

### DLPFC best distinguishes control-type at go cue, while M1 is best at target acquisition

To gain further insight into the differences between neural representations of BMI control and manual control, we used a *2-class* LDA model to distinguish between BMI control and manual control only (Figure 1c). At the go cue, using power features from DLPFC resulted in significantly higher classifier performance than using power features from M1 or Cd (Figure 4b-c, Supplementary Figure 6a). Additionally, the classification accuracy of models using power features from either DLPFC + Cd or M1 + DLPFC was not significantly different from that of models using power features from all ROIs for either subject. At target acquisition, models using power features from M1 resulted in significantly higher classifier performance than the classifiers using power features from DLPFC or Cd for Monkey H and from Cd for Monkey Y (Figure 4d-e, Supplementary Figure 6b). Additionally, the classification accuracy of models using power features from either M1 + DLPFC or M1 + Cd was not significantly different from that of models using power features from all ROIs for either subject. To ensure that these results were consistent across BMI learning, both the go cue and target acquisition results were similar across BMI learning stages (Supplementary Figure 7). These results suggest that most of the information for distinguishing between BMI and manual control at the go cue is present in DLPFC, while most of the information at target acquisition is present in M1. This difference may indicate that cognitive mechanisms known to rely more heavily on DLPFC or M1, such as action-planning or movement-execution, are more involved at the go cue and at target acquisition, respectively.

Though DLPFC and M1 hold key information for distinguishing BMI and manual control at the go cue and at target acquisition, respectively, models using power features from all individual ROIs at both task events yielded above-chance classification accuracy (Figure 4b and 4d). This indicates that each individual ROI holds important information for the *2-class* model. Additionally, models with features from all 3 ROIs combined consistently yielded the highest classification accuracy (Supplementary Figure 6). This increase in accuracy obtained by combining information across ROIs indicates that there is non-overlapping information encoded by each ROI.

### Information flow to M1 during both BMI and manual control

To investigate how these regions interact with one another during BMI and manual control, we evaluated the directed functional connectivity, or effective connectivity, using Granger causality. Granger causality is a statistical test for determining whether one time-series is useful in predicting another, and therefore provides insight into the direction of information flow between these ROIs. For successful trials under BMI and manual control, we calculated Granger causality in the 500 ms LFP segments following the go cue and those preceding target acquisition. The same calculations were also performed on random non-overlapping 500ms windows from the baseline period. At both task events, we found significantly above-chance effective connectivity in every ROI→ROI interaction direction during all task types: BMI control, manual control, and baseline (see Methods for details). This indicates that every ROI sends information to every other ROI. Because all ROIs exhibit significant reciprocal interactions, even during baseline, we isolated task-specific directed information flow by calculating a normalized Granger causality that compared interactions during BMI and manual control to interactions during baseline. A normalized Granger causality value greater than zero indicates an increase from baseline, while a value less than zero indicates a decrease from baseline. This normalized value provides insight into which interactions are task-specific and accounts for variability in baseline information flow across days. We first analyzed this normalized metric of directed information flow during BMI control. We observed an increase in normalized Granger causality in the DLPFC→M1 and Cd→M1 directions and a decrease in the M1→Cd direction in both monkeys at both the go cue and target acquisition (Figure 5a-b). Additionally, there was an increase in the DLPFC→Cd direction for Monkey H and a decrease in the Cd→DLPFC direction for Monkey Y at the go cue (see Supplementary Table 3). These results suggest information flow from DLPFC→M1 and from Cd→M1 during both task events, as well as information flow from Cd→DLPFC at the go cue. Next, we determined the change in directed information flow from baseline during manual control. We observed a similar increase in normalized Granger causality from DLPFC→M1 and from Cd→M1 in both monkeys at both task events (Figure 5a-b). However, unlike during BMI control, DLPFC ↔ Cd interactions were inconsistent across animals. While there were significant differences between BMI and manual control normalized Granger causality within a subset of interaction directions, these differences were inconsistent across monkeys and mainly differed in amplitude rather than direction (Figure 5a-b, see Supplementary Table 4). Overall, these results suggest information flow from DLPFC→M1 and Cd→M1 is present during both BMI and manual control.

For both BMI and manual control, most reciprocal interactions underwent opposite changes from baseline. To consolidate these results, we calculated the net normalized Granger causality by taking the difference between each pair of reciprocal directed interactions (Figure 5c). This metric, which represents the overall predominant direction of task-specific information flow, provides a clear representation of the network between our ROIs. We refer to information flow between two ROIs as significantly directed if we found the difference in normalized Granger causality to be significantly different from zero. During BMI control, we found that information was significantly directed from DLPFC→M1 and Cd→M1 for both monkeys at both task events (Supplementary Tables 3a, 3b). At the go cue, information was also significantly directed from Cd→DLPFC. Just as with our control-type classification results, we assessed whether this directed network was present across BMI learning stages and found no differences (Supplementary Figure 8). Under manual control, we found similar directions of net change in information flow from baseline. The net normalized Granger causality was significantly directed, from DLPFC→M1 for both monkeys across both task events (Supplementary Tables 3a, 3b), similar to BMI control (Figure 5f-g). The interaction from Cd→M1 was significantly directed for Monkey Y, but was not above chance for Monkey H. Interactions between DLPFC and Cd were significantly directed from DLPFC→Cd in Monkey H, rather than Cd→DLPFC as observed during BMI control, but were not significant for Monkey Y. Within each monkey, these results were consistent across task events. Overall, there were many similarities in the net direction of information flow across both BMI and manual control. Under both control types, the net direction of Granger causality increased from DLPFC→M1 relative to baseline at both the go cue and target acquisition. However, the net direction of information flow between Cd and M1 and between DLPFC and Cd was not consistent across monkeys during manual control. Therefore, DLPFC may play an upstream role for both BMI and manual control, while the upstream role of Cd may be more clear for BMI control.

Our results demonstrate the presence of distinct neural representations of BMI and manual control in M1, DLPFC, and Cd. More specifically, we found that neural activity from DLPFC and M1 best distinguish between control types, at the go cue and target acquisition, respectively. Further, we identified net directed information flow from DLPFC→M1 in both BMI and manual control, and from Cd→M1 in BMI control. Together, these results suggest that DLPFC plays an important upstream role in both BMI and manual control tasks, with especially distinct neural representations during a period in the task that requires action planning, selection, and initiation.

## Discussion

In this study, single- and multi-unit motor cortical recordings from a semi-chronic microdrive array were used to successfully control a BMI. We leveraged simultaneous LFP recordings from M1, DLPFC, and Cd to investigate differences in distributed activity within cortical and subcortical networks during motor BMI control and during overt motor (manual) control. Our results demonstrate the presence of distinct neural representations of BMI and manual control in M1, DLPFC, and Cd. Notably, we found that neural activity from DLPFC and M1 best distinguishes control types at the go cue and at target acquisition, respectively. Further, we identified net directed information flow from DLPFC→M1 in both BMI and manual control and from Cd→M1 in BMI control. Together, these results suggest that DLPFC plays an important upstream role in both BMI and manual control tasks, with especially distinct neural representations during a period in the task that requires action planning, selection, and initiation. On the contrary, M1 is a net-receiver of information and is better at distinguishing control types during a period in the task that requires more fine effector control. Overall, this work provides evidence of coordinated network activity between M1, DLPFC, and Cd during BMI control that is similar yet distinct from manual control.

We compared spectral power during baseline, BMI control, and manual control and found that theta power in M1, DLPFC, and Cd was significantly higher during the control tasks than during baseline at the go cue. In contrast, there were no significant distinctions in theta power at target acquisition. Previous work has demonstrated the involvement of theta activity in goal-directed information processing [39], [40], with increased theta activity in the prefrontal cortex observed at decision points where choice-relevant information is under consideration. In the center-out task, the go cue can be thought of as a decision point, where subjects must choose a cursor trajectory, while the target acquisition period involves less choice-relevant information processing. Therefore, this observed increase in theta power across all ROIs at the go cue may be related to trajectory selection and planning. In contrast to theta power, beta power in M1, DLPFC, and Cd at the go cue was significantly lower during the control tasks than at baseline. At target acquisition, only M1 beta power remained distinct between baseline and the two control tasks. Beta power in the prefrontal cortex has been implicated in attention and task-specific rule representation [41], [42]. Beta power in the motor cortex and striatum can be modulated by task-relevant cues, movement preparation, and movement execution [43]–[46]. Therefore, beta power within each of our ROIs may play an important role in maintaining task-relevant information, with DLPFC and Cd beta power modulated more specifically during trajectory planning at the go cue. These differences in oscillatory activity in the theta and beta frequency bands between baseline and the two control tasks suggest that this activity, which has previously been observed during planning, movement, and other goal-directed tasks, may be utilized with similar functionality in both BMI and manual control.

We found distinct BMI and manual neural representations in M1, DLPFC, and Cd at both task events when comparing combinations of frequency bands. We also found that control-type classification accuracy always increased with the addition of neural activity from a second or third ROI, suggesting that each ROI holds some non-overlapping information for differentiating between BMI and manual control. One major difference between our BMI and manual control tasks is the lack of proprioceptive feedback in the BMI task. Many models of how the motor cortex controls movement incorporate sensory feedback as a relevant factor [47]. One potential explanation for observed differences in M1 activity is the difference in sensory feedback between the two control types. The differences between control types in M1 LFP may also be related to changes in the properties of individual M1 neuron activity. Distinct changes in neural activity between BMI and manual control have been previously observed in single- and multi-unit motor cortical activity, in the form of changes in preferred direction and modulation depth [29], [48]–[51]. The differences we observed between BMI control and manual control in M1 LFP may reflect or drive these distinct single- and multi-unit properties across control types. DLPFC activity also successfully differentiated between BMI and manual control and is largely involved in action planning, selection, and initiation [13], [18], [26]. Thus, the distinct neural representations in DLPFC activity may reflect differences between action planning, selection, and initiation between BMI and manual control. Furthermore, we also found that DLPFC activity was better than other ROIs at distinguishing control types at the go cue, a portion of the task that may rely more heavily on action planning, selection, and initiation, further supporting the hypothesis that the differences in DLPFC activity are related to differences in these cognitive processes. Additionally, the prefrontal cortex has been implicated in rule representations and in task- and rule-switching [18], as has the basal ganglia [52], [53]. In our experiment, proficient motor BMI and manual control could be viewed as two rules dictating how the subjects must perform the center-out task. Thus, the distinct neural representations in DLPFC and Cd may be involved in differentiating between these two control types as distinct rules.

Behavioral performance is another major difference between BMI and manual control, with BMI behavioral performance significantly changing across days while manual performance remains more stereotyped and consistent. However, despite these changes in the BMI control task, our classification results did not differ across learning stages, implying that the distinctions in neural representations do not reflect skill learning and changes in performance. In contrast, previous literature has shown decreases in DLPFC activity and increases in striatal activity across motor learning [54]. Additionally, decreased PFC activity has been observed across learning of a 1-D BMI task in humans [8], [35] as well as an increase in cortico-striatal coherence across learning of a 1-D BMI task in rodents [32]. However, the 2-D, center-out BMI task used in this study is more challenging than both an overtrained overt motor control task and a 1-D BMI control task. Because the task used in this study required the animal to hold a center target in order to successfully initiate a trial, even trial initiation required some amount of proficient control from each NHP. Thus, any truly initial learning that would need to occur prior to this baseline level of proficiency would not have been captured by our experimental design. Although our results were consistent across early and late learning stages, it is important to note that even at their most proficient BMI control, both subjects performed the task more slowly under BMI control and never achieved the same target acquisition time as manual control. Thus, even proficient BMI control may involve greater online error correction than manual control. Previous studies have discussed the role of the striatum in error-based motor learning and encoding error in movement observation [55]–[58]. The distinctions we observe in Cd may represent this difference in online error correction across control types. However, the exact role of Cd in error correction is still debated and comparing the role of other regions implicated in error prediction, such as the cerebellum, is necessary [59]. While further research is necessary to understand which cognitive processes are relevant to each brain region in BMI control, our results expand on prior knowledge of distinct neural representations between BMI and manual control in M1 and also show distinctions in the neural activity of distributed brain regions beyond M1 including in DLPFC and Cd.

To investigate how these regions interact with one another during both BMI and manual control, we quantified directed functional connectivity using Granger causality. We observed net directed information flow from DLPFC→M1 during both BMI and manual control. Previous studies have identified structural and functional connections between prefrontal and motor cortices, with the prefrontal cortex sending information to the motor cortex regarding both motor regulatory functions [13], [60] and goal-directed behaviors [25]. Our results support this upstream information processing model during overt motor control and suggest that a similar model is present during BMI control. The direction of net information flow was consistent across the go cue and target acquisition, suggesting that the information-processing network is stable despite shifts in primary cognitive demands throughout an individual trial and may be representative of a more general state of goal-directed behavior.

We also observed net directed information flow from Cd→M1 during BMI control. Much of the previous literature on cortico-striatal connections has focused on motor execution directed from M1 to the striatum due to the existence of direct structural connections between them [20]. However, recent studies in mice have confirmed the presence of a cortico-striatal thalamo-cortical feedback loop that is relevant in motor control, suggesting that indirect modulation of the motor cortex by the striatum also plays an important role in motor control [22], [61]–[63]. Additionally, a recent study in humans suggests that basal ganglia may contribute to the regulation of sensorimotor cortical regions [64]. While the bottom-up indirect feedback from the striatum to the motor cortex has not received as much attention, particularly in NHPs, our results suggest that this directed functional connection may be an important area of future study in BMI control.

Altogether, this study leverages simultaneous cortical and subcortical recordings during the same task under BMI control and under overt motor control to identify control-specific neural representations within M1, DLPFC, and Cd and to uncover an information-processing network from DLPFC and Cd to M1 during BMI control. Our results contribute to a growing body of work elucidating the neural mechanisms underlying BMI control, improving our understanding of BMIs as a scientific tool and as a therapeutic device. This study provides insight into the cortical and subcortical circuits supporting both motor and high-level cognitive functions associated with BMI control.

## Methods

### Surgery

Two rhesus macaques were implanted with recording chambers. Chamber positions were calculated based on images obtained from 3T magnetic resonance imaging (MRI) (Siemens Medical Solutions, Malvern, PA) scans of each subject’s brains. Regions of interest, including primary motor cortex (M1), dorsolateral prefrontal cortex (DLPFC), and caudate (Cd), were manually traced in 3D Slicer [65] using the Paxinos primate atlas as a reference [66]. The resulting neuroanatomical models were used to decide on stereotaxic coordinates for implantation. All procedures were conducted in compliance with the National Institute of Health (NIH) Guide for the Care and Use of Laboratory Animals and were approved by the University of California at Berkeley Institutional Animal Care and Use Committee.

### Large-Scale Semi-Chronic Microdrive

We used custom large-scale semi-chronic microdrive arrays for recording in both animals (Gray Matter Research, MT). These arrays allowed us to simultaneously record from regions of interest at different depths using independently-moveable single microelectrodes (n = 124). Electrodes consisted of both glass-coated Tungsten electrodes (Alpha Omega, Nof HaGalil (Nazareth Illit), Israel) and Platinum-Iridium electrodes (MicroProbes for Life Sciences, Gaithersburg, MD). Throughout implantation and recovery, electrodes were stored inside of the microdrive. The impedance of the electrodes was monitored using a TDT NanoZ (Tucker-Davis Technologies, Alachua, Florida) while advancing them into the brain. First, electrodes were lowered out of the microdrive until they penetrated the dura. While most electrodes successfully entered the brain (Monkey H: 79/124; Monkey Y: 64/124), a subset broke upon penetrating the dura (Monkey H: 45/124; Monkey Y: 60/124). These recording channels were excluded from all analyses. Electrodes that successfully entered the brain were advanced until their respective neural targets were reached and unit activity was detected. The estimated target depth of each electrode was calculated using the neuroanatomical models built in 3D Slicer. Ultimately, we successfully lowered a subset of electrodes into each region of interest (Monkey H: M1 = 27, PMd = 10, DLPFC = 22, Cd = 8; Monkey Y: M1 = 12, PMd = 12, DLPFC = 16, Cd = 8).

### Intracortical Recording

Neural data were recorded using the OmniPlex Neural Recording Data Acquisition System (Plexon Inc, Dallas, TX). Single- and multi-unit activity was sorted prior to beginning recording sessions using an online sorting application (Sort Client, Plexon Inc, Dallas, TX). Wideband activity was recorded at 5 kHz. LFP activity was obtained by low-pass filtering at 250 Hz, notch filtering at 60 Hz and 120 Hz, and down-sampling to 1 kHz. LFP activity was common median referenced by first z-scoring activity within each channel (*x*_*i*_ *where i*: {1, 2, … *n*}) by subtracting the mean and dividing by the standard deviation in each recording session (*s*)

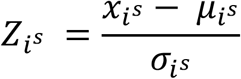

and then subtracting the median value across all channels at each time point (*t*).

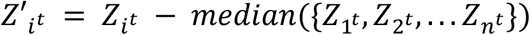

After subtracting the median to remove common signals, likely low frequency noise, across channels, the LFP activity was multiplied by the standard deviation and the mean was added to restore the LFP to its original scaling.

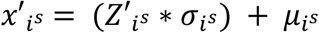

Common median referencing was used instead of common average referencing to avoid influence from outliers. The common median referenced LFP activity was then z-scored using the mean and standard deviation within channel across all recording sessions within a day (1, 2, … *S*).

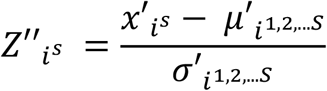

### Center-Out Task

Subjects performed a self-paced, center-out, reaching task to eight targets. Trials were initiated by moving a cursor to the central target. A successful trial required a short hold at the center, moving to the peripheral target within a specified time, and a brief hold at the target. Successful trials resulted in a juice reward; failed trials were repeated up to 10 times before a new target was presented. Target directions were presented in a pseudo-randomized order.

Subjects were first overtrained in the center-out task performed with arm movements before starting BMI. In this manual control (MC) version of the task, the subject’s arm moved in a KINARM exoskeleton (BKIN Technologies, Kingston, ON, Canada) that restricted movements to the horizontal plane. Neural activity recorded during MC was used to train a BMI decoder. Using this BMI decoder, the animals performed the same task under BMI control (BC). During BC, the animals’ arms were restricted in a fixed position within the exoskeleton and the animals were required to move the cursor to the target by modulation of motor cortex activity.

### Brain-Machine Interface

Subjects learned to control a two-dimensional BMI cursor in real-time using a fixed velocity Kalman Filter (KF) decoder [67]–[69]. The KF assumes two linear models:

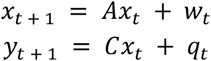

where and are the cursor state and neural activity at time t, respectively. The first equation represents the state-transition model, which describes the state of the cursor over time. It is specified by the state-transition matrix and additive Gaussian noise term. Equation 2 represents the observation model and describes the relationship between neural activity and cursor state. It is parameterized by the observation matrix and additive Gaussian noise. Neural activity was input as a vector of spike counts in 100 ms bins from the selected direct units.

Decoder parameters were initialized from neural and cursor kinematic data collected during the MC version of the task at the beginning of each recording day. Maximum likelihood estimation methods were used to fit initial parameters. Neural data from 10-20 single- and multi-units recorded from motor cortex were selected for BMI control each day (i.e., direct units). The population of direct units was highly overlapping from day to day, but there was some variability as units dropped out or new ones appeared. For Monkey Y, closed-loop decoder adaptation (CLDA) was performed using the SmoothBatch algorithm before fixing the BMI decoder. This algorithm uses knowledge of task goals (i.e., reaching targets) to infer a subject’s intent. The intended kinematics and observed neural activity during closed-loop BMI were used to re-estimate KF parameters. The SmoothBatch algorithm [70]–[72] re-estimates the observation model of the KF (matrices C and Q), and updates were constrained to enforce smoothness. CLDA was typically run for 2-5 minutes to provide the subject with adequate performance to allow successful reaches to all targets. For Monkey H, the initial decoder trained from MC data was used.

### Neural Data Analysis

All analyses were performed in Python with custom-written routines utilizing publicly available software packages including scipy [73], numpy [74], sklearn [75], and statsmodels [76]. Unless otherwise specified, analyses involving activity from DLPFC and Cd were performed on the average LFP signal across all channels within DLPFC and Cd, respectively. Analyses involving M1 were performed on the average LFP signal across all channels that were used to record single- or multi-unit activity that was input to the BMI decoder.

### Linear Classifiers

In order to understand which of our ROIs were most predictive of control type, we trained linear classifiers to predict between control types using power from different ROIs and frequency bands as input features. We used linear discriminant analysis (LDA) models using singular value decomposition (SVD) from the sklearn toolbox in Python. Power in the theta (4-8 Hz), alpha (8-13 Hz), beta (13-35 Hz), gamma (35-75 Hz), and highgamma (75-150 Hz) frequency bands was calculated using Welch’s method on segments of LFP recorded during each control type. For BMI and manual control, power was computed in the 500 ms window following the go cue and in the 500 ms window preceding target acquisition on successful trials. Power estimates were averaged across all channels in each ROI prior to being used as input features. The same calculations were also performed in random non-overlapping 500ms windows during the baseline period. To avoid negative effects on imbalanced data between classes, the number of trials selected for each task type was matched to the minimum number of trials per individual task type each day by selecting a random subset of trials from the larger class to match that minimum. The corresponding number of windows were selected from the baseline period as well.

Results for each model were validated using a 10-fold cross-validation within each recording day. To validate whether classification accuracy results were significantly above chance, we used a paired t-test to compare each result to a chance accuracy. For each frequency band and ROI combination, chance accuracy was calculated by randomly shuffling class labels and rerunning the LDA classification, trained on 90% of data and tested on 10% of data, 1000 times for each day.

To determine whether particular frequency bands were useful for distinguishing between task types, we computed the LDA classification accuracy using each individual frequency band from each ROI as well as using all frequency bands from each ROI as input (Supplementary Figure 3). We determined that all frequency bands in all ROIs could predict the task type significantly above chance and that the classifier trained on power features from all 5 frequency bands outperformed the classifiers using individual bands, so all subsequent analyses use classifiers trained on all 5 frequency bands from each ROI.

### Granger Causality

Granger causality was used to estimate the directional functional connectivity among all pairs of regions of interest. Granger causality relies on an autoregressive (AR) modeling framework, in which future values of a time series are modeled as a weighted combination of past values of time series. The quality of an AR-model is assessed by quantifying the variance of the model’s residuals. If the variance of the AR-model’s residuals is reduced by the inclusion of past measurements from a second time series, then the second time series is said to Granger-cause or G-cause the first [77], [78]. Applying this logic, we obtained Granger causality estimates using simultaneously recorded LFP signals. We compared Bayesian information criterion (BIC) of AR-models of different orders, p, for each combination of LFP signals obtained from all pairs of regions during each trial from all recording sessions. We selected p=13 to minimize BIC in the average case, allowing for the signals to be sufficiently long enough to capture the data structure without over-parameterization.

For successful trials under BMI and manual control, we calculated Granger causality in the 500 ms average LFP segments following the go cue and those preceding target acquisition. The same calculations were also performed on random non-overlapping 500ms average LFP segments from the baseline period. To obtain an estimate of spurious interactions, we calculated Granger causality between the average LFP during baseline and the average LFP during BMI control. Because these signals were recorded at separate times in different tasks, any interactions that occur between regions would be artifactual. True interactions between regions within baseline and BMI control were considered significant if they were statistically different from this null distribution. The average Granger causality value across BMI control trials within each day was determined and compared to that of the null distribution. A null distribution for manual trials was calculated and compared using the same protocol.

We compared ROI→ROI Granger causality (*G*_*Y*→*X*_) in each interaction direction to the null distribution via an unpaired t-test and found that every interaction was significantly greater than the null distribution during all task types: BMI control, manual control, and baseline. In order to isolate task-relevant interactions, we computed the normalized Granger causality for BMI and manual control as follows:

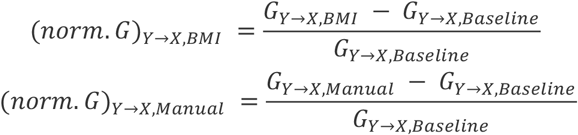

Here, a normalized Granger causality greater than zero indicates an increase from baseline, whereas a normalized Granger causality less than zero indicates a decrease from baseline. To further assess the direction of information flow, we also calculated the net normalized Granger causality by subtracting the value of normalized Granger causality between ROI → ROI reciprocal interactions.

### Quantification & Statistical Analyses

All analyses were performed within a single recording day and error is depicted across days. For analyses comparing two distributions or comparing a single distribution to a value, two-sample or one-sample t-tests were used, respectively. Bonferroni correction was used post hoc to correct for multiple comparisons. Significance is reported after correction for multiple comparisons.

### Ethical approval

All procedures were conducted in compliance with the NIH Guide for the Care and Use of Laboratory Animals and reporting in the manuscript follows the recommendations in the ARRIVE guidelines. All procedures were approved by the University of California at Berkeley Institutional Animal Care and Use Committee under protocol ID AUP-2014-09-6720-2.

## Acknowledgements

We thank Maki Kitano for assistance with NHP care, handling, and training, Paul Botros for methodological advice, and Charles Gray for help with surgical procedures.This work was supported by the National Science Foundation Graduate Research Fellowship (to E.L.Z.), the National Institute of Mental Health Award R01MH117763 (to J.D.W.), and the National Institute of Health Award R01NS106094 (to J.M.C.).

## Author Contributions

E.L.Z., G.F.S., J.D.W., and J.M.C. designed and performed research. E.L.Z. and G.F.S. analyzed the data. E.L.Z., G.F.S., and N.V-L. interpreted the data. E.L.Z. and G.F.S. wrote the manuscript text and prepared the manuscript figures. N.V-L., J.D.W., and J.M.C edited the manuscript. All authors reviewed the manuscript.

## Data Availability Statement

The datasets analyzed during the current study will be publicly available on Figshare upon publication.

