## Supplemental Information for "Distinct neural representations during a brain-machine interface and manual reaching task in motor cortex, prefrontal cortex, and striatum"

**Supplementary Information**

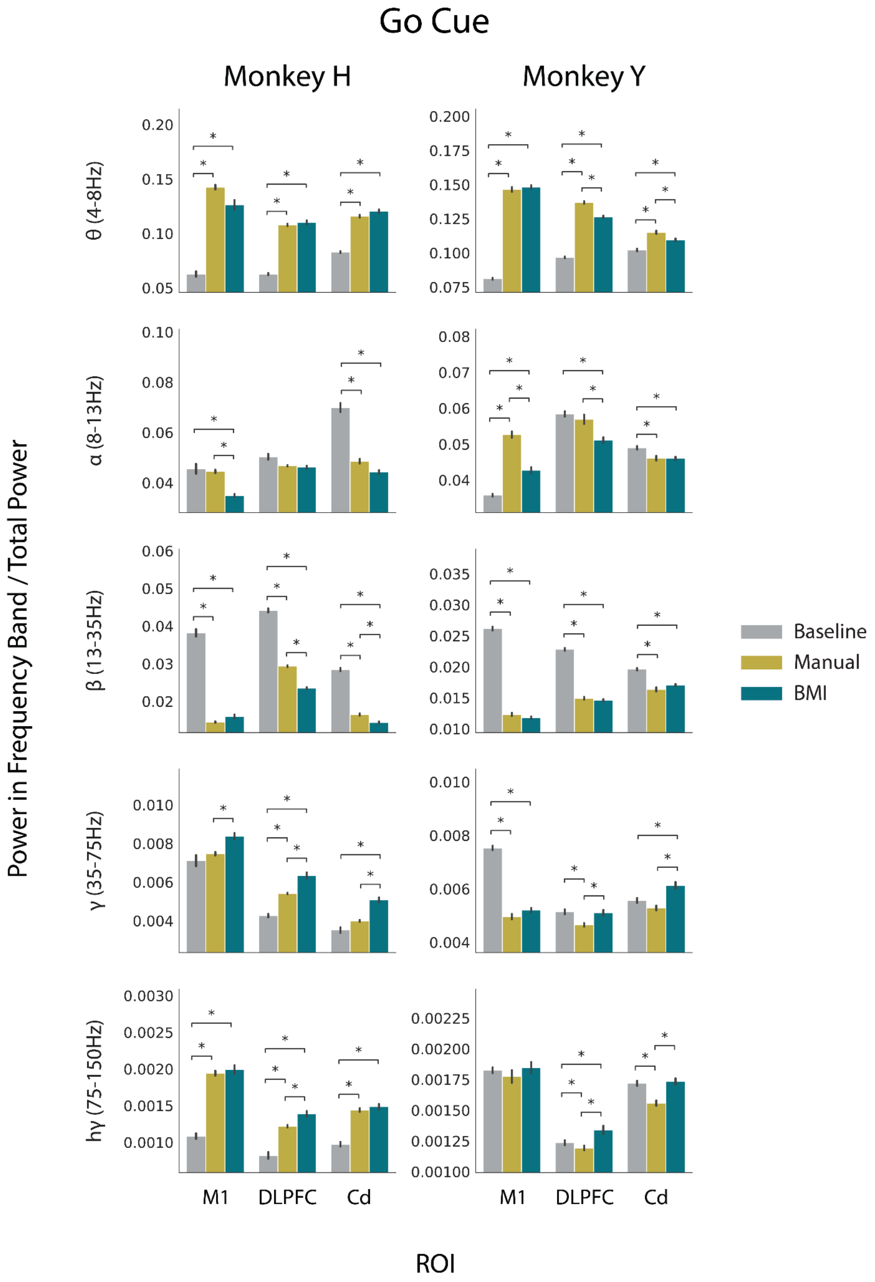


**Supplementary Figure 1. Spectral power in M1, DLPFC, and Cd during baseline, manual control, and BMI control at the go cue**. LFP was decomposed into 5 distinct frequency bands. The average power normalized to total power in each of these bands in each ROI was compared during baseline (gray), manual (yellow), and BMI (teal) at the go cue for Monkey H (left) and Monkey Y (right). Error bars represent standard error mean across days. Normalized spectral power estimates significantly differing between task types after Bonferroni correction for multiple comparisons are indicated with an asterisk.

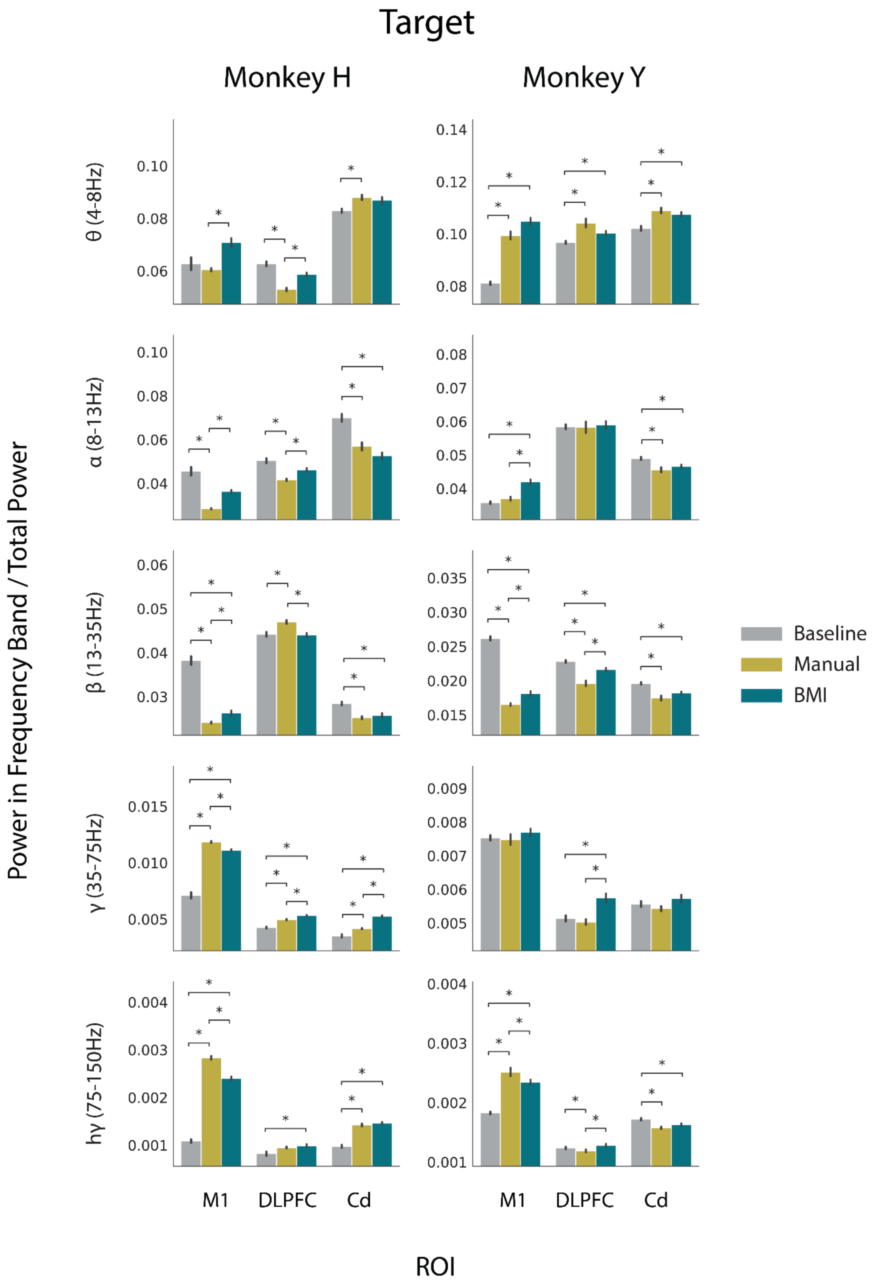


**Supplementary Figure 2. Spectral power in M1, DLPFC, and Cd during baseline, manual control, and BMI control at target acquisition**. LFP was decomposed into 5 distinct frequency bands. The average power normalized to total power in each of these bands in each ROI was compared during baseline (gray), manual (yellow), and BMI (teal) at target acquisition Monkey H (left) and Monkey Y (right). Error bars represent standard error mean across days. Normalized spectral power estimates significantly differing between task types after Bonferroni correction for multiple comparisons are indicated with an asterisk.

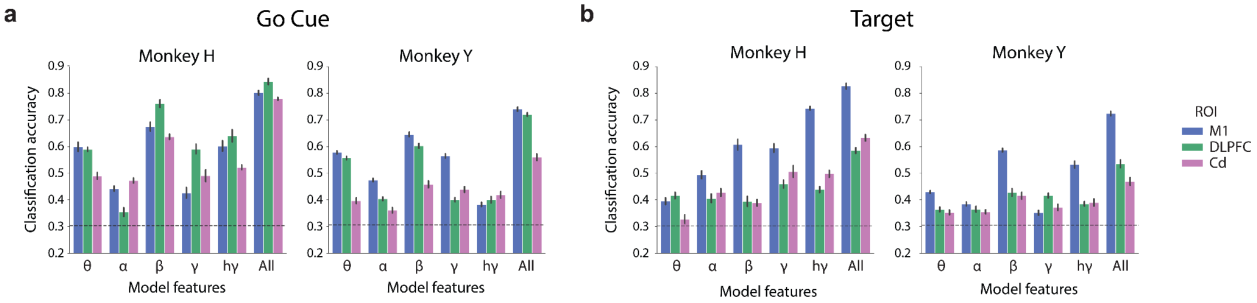


**Supplementary Figure 3. Task-type classification accuracy is above chance for all individual frequency bands and highest for the combination of all frequency bands in all ROIs.** (a) Mean 10-fold cross-validated classification accuracy across days for a *3-class* LDA classifier trained to distinguish between BMI control, manual control, and baseline using individual frequency bands and all combined bands at the go cue for Monkey H (left) and Monkey Y (right). Chance accuracy shown as a dashed line. Error bars represent standard error of the mean across days. (b) Same as (a) but for target acquisition, rather than go cue.

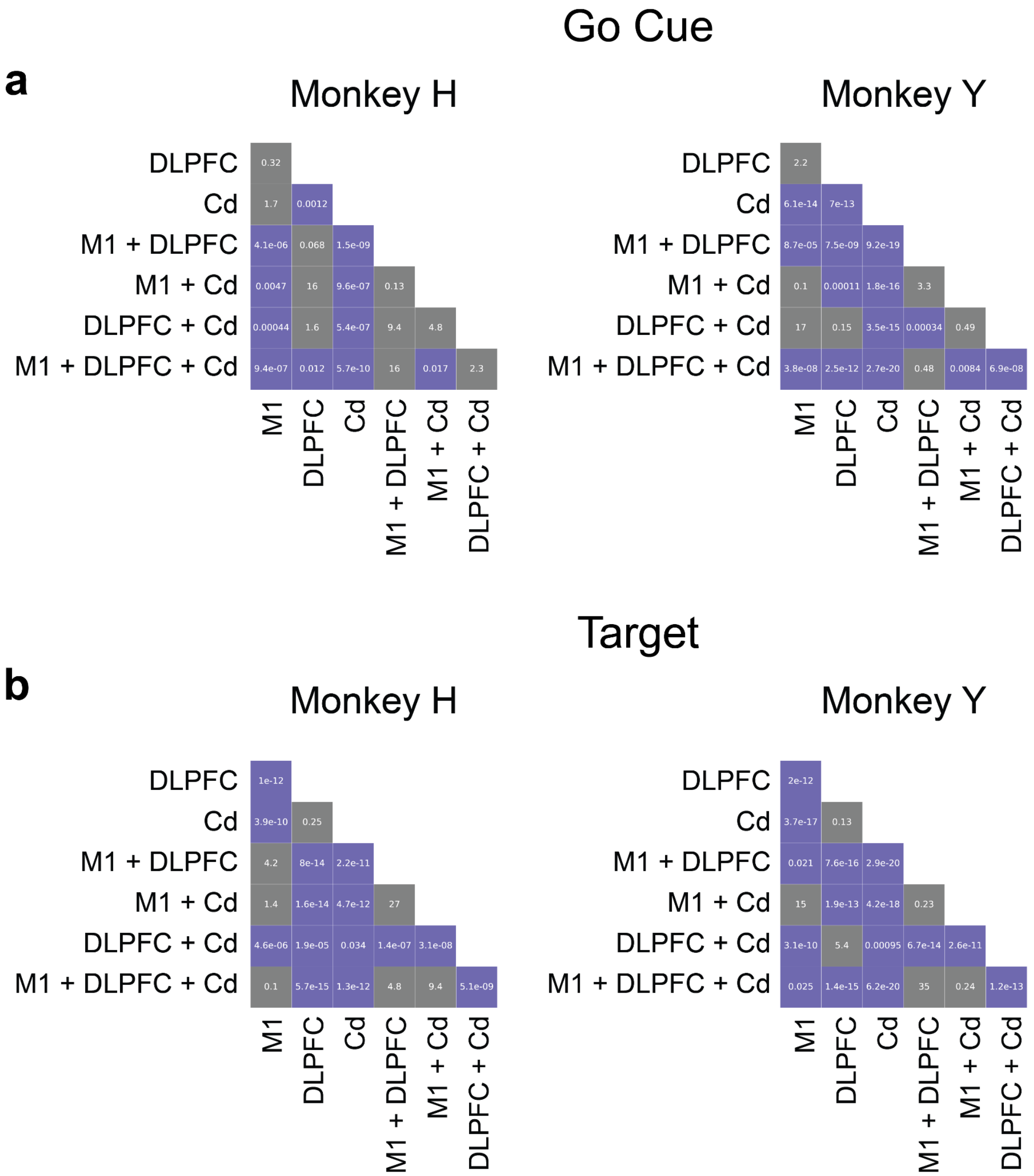


**Supplementary Figure 4. Statistical significance values for pairwise comparisons of all *3-class* LDA models.** (a) Bonferroni corrected p-values for all unpaired t-tests comparing cross-validated classification accuracy obtained from LDA classifiers with input from different combinations of ROIs at the go cue for Monkey H (left) and Monkey Y (right). Purple squares indicate a Bonferroni corrected p-value < 0.05. Gray squares indicate a Bonferroni corrected p-value > 0.05. (b) Same as (a) but for features obtained at target acquisition, rather than at go cue.

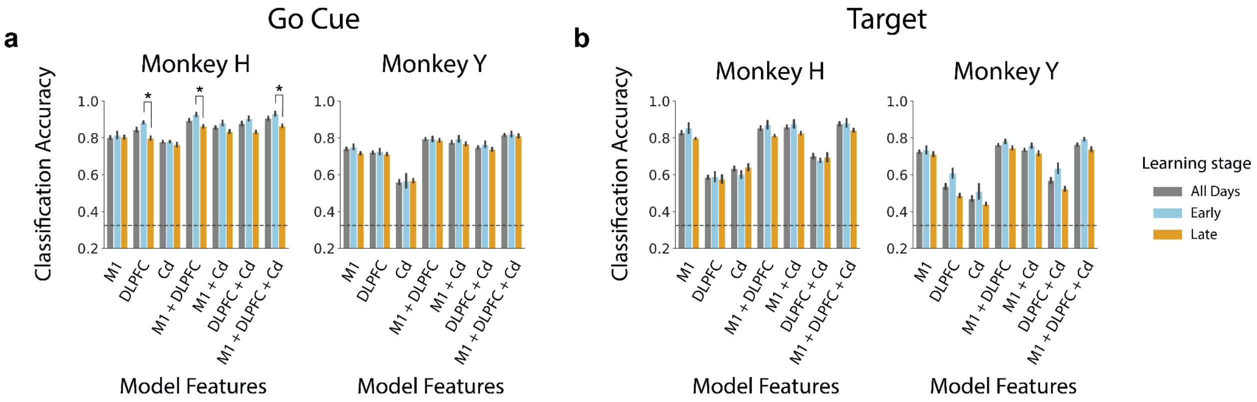


**Supplementary Figure 5. Classification accuracy using the *3-class* LDA model is similar across BMI learning stages.** (a) Mean 10-fold cross-validated classification accuracy across all days (gray), early learning days (blue), and late learning days (orange) for a *3-class* LDA classifier trained to distinguish between BMI control, manual control, and baseline using all frequency bands from individual ROIs or combinations of ROIs at the go cue for Monkey H (left) and Monkey Y (right). Chance accuracy shown as a dashed line. Error bars represent standard error mean across days. (b) Same as (a) but for target acquisition, rather than go cue.

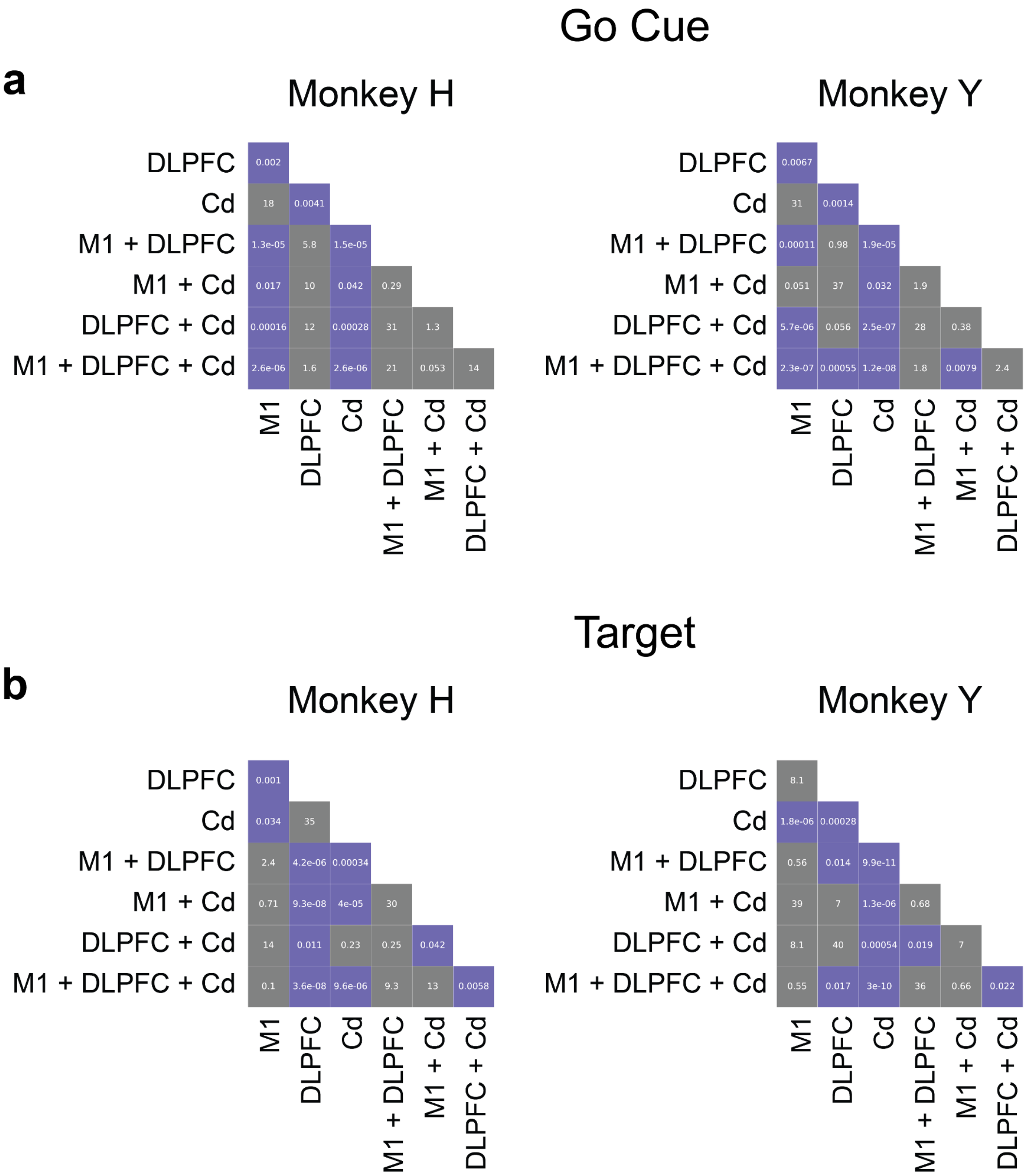


**Supplementary Figure 6. Statistical significance values for pairwise comparisons of all *2-class* LDA models.** (a) Bonferroni corrected p-values for all unpaired t-tests comparing cross-validated classification accuracy obtained from LDA classifiers with input from different combinations of ROIs at the go cue for Monkey H (left) and Monkey Y (right). Purple squares indicate a Bonferroni corrected p-value < 0.05. Gray squares indicate a Bonferroni corrected p-value > 0.05. (b) Same as (a) but for features obtained at target acquisition, rather than at go cue.

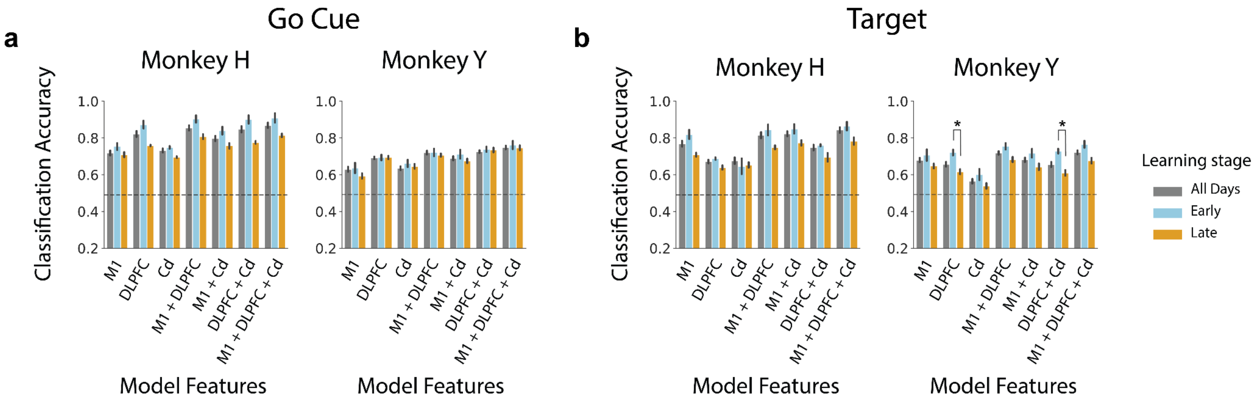


**Supplementary Figure 7. Classification accuracy using the *2-class* LDA model is similar across BMI learning stages.** (a) *Mean 10-fold cross-validated classification accuracy across all days (gray), early learning days (blue), and late learning days (orange) for a 2-class LDA classifier trained to distinguish between BMI and manual control using all frequency bands from individual ROIs or combinations of ROIs at the go cue for Monkey H (left) and Monkey Y (right). Chance accuracy shown as a dashed line. Error bars represent standard error mean across days. (b) Same as (a) but for target acquisition, rather than go cue.*

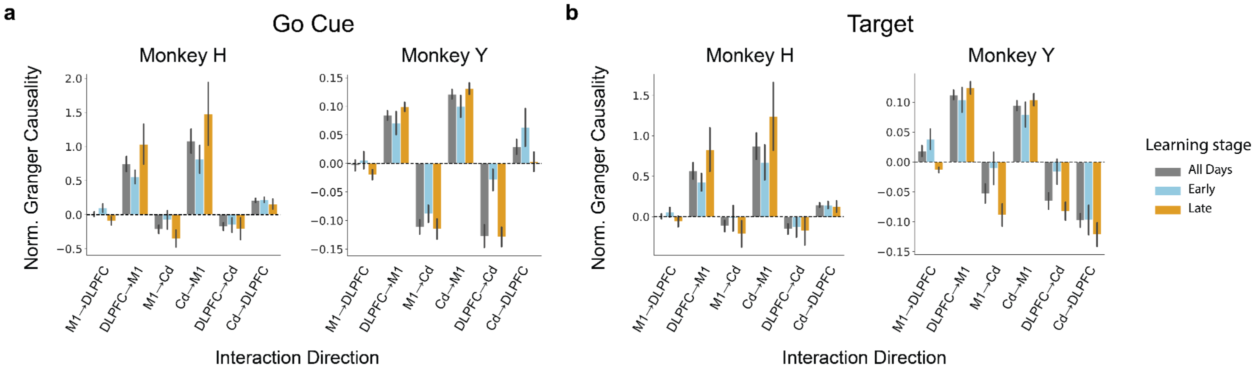


**Supplementary Figure 8. Normalized granger causality is similar across BMI learning stages** (a) Normalized Granger causality estimates at the go cue during BMI control across all days (gray), early BMI learning (blue), or late BMI learning (orange). Error bars represent standard error mean across days. (b) Same as (a) but for target acquisition, rather than go cue.

**Go Cue:** Comparing Average Power Between Task Types

| Subject | Frequency Band | Comparison | M1 t-statistic | M1 Bonferroni corrected p | DLPFC t-statistic | DLPFC Bonferroni corrected p | Cd t-statistic | Cd Bonferroni corrected p |
|---|---|---|---|---|---|---|---|---|
| Monkey H | Theta (4-8Hz) | Baseline vs Manual | **-24.496** | **5.816e-10*** | **-31.148** | **3.393e-11*** | **-29.790** | **5.758e-11*** |
| | | Baseline vs BMI | **-12.170** | **1.860e-06*** | **-23.000** | **1.221e-09*** | **-20.825** | **3.913e-09*** |
| | | Manual vs BMI | 3.816 | 0.110532 | -1.461 | 7.633671 | -3.380 | 0.246192 |
| | Alpha (8-13Hz) | Baseline vs Manual | 0.507 | 27.953291 | 3.006 | 0.492121 | **11.515** | **3.441e-06*** |
| | | Baseline vs BMI | **5.242** | **9.317e-03*** | 3.798 | 0.114247 | **13.665** | **5.055e-07*** |
| | | Manual vs BMI | **11.801** | **2.622e-06*** | 0.835 | 18.898263 | 3.964 | 0.084514 |
| | Beta (13-35Hz) | Baseline vs Manual | **24.787** | **5.059e-10*** | **23.012** | **1.213e-09*** | **29.715** | **5.933e-11*** |
| | | Baseline vs BMI | **19.604** | **7.927e-09*** | **34.524** | **9.998e-12*** | **37.511** | **3.725e-12*** |
| | | Manual vs BMI | -3.343 | 0.263690 | **24.101** | **7.044e-10*** | **12.016** | **2.145e-06*** |
| | Gamma (35-75Hz) | Baseline vs Manual | -1.275 | 10.186384 | **-12.249** | **1.730e-06*** | -3.566 | 0.174739 |
| | | Baseline vs BMI | -3.511 | 0.193216 | **-13.590** | **5.376e-07*** | **-9.780** | **2.049e-05*** |
| | | Manual vs BMI | **-5.283** | **8.719e-03*** | **-6.492** | **1.337e-03*** | **-10.371** | **1.085e-05*** |
| | Highgamma (75-150Hz) | Baseline vs Manual | **-19.798** | **7.067e-09*** | **-9.234** | **3.785e-05*** | **-19.968** | **6.396e-09*** |
| | | Baseline vs BMI | **-15.355** | **1.338e-07*** | **-14.554** | **2.469e-07*** | **-17.135** | **3.785e-08*** |
| | | Manual vs BMI | -0.778 | 20.321279 | **-4.749** | **0.021287*** | -1.599 | 6.109760 |
| Monkey Y | Theta (4-8Hz) | Baseline vs Manual | **-46.227** | **5.844e-21*** | **-57.829** | **5.493e-23*** | **-8.890** | **6.564e-07*** |
| | | Baseline vs BMI | **-58.992** | **3.624e-23*** | **-33.419** | **4.865e-18*** | **-8.840** | **7.207e-07*** |
| | | Manual vs BMI | -1.215 | 10.708290 | **12.666** | **1.203e-09*** | **4.909** | **3.345e-03*** |
| | Alpha (8-13Hz) | Baseline vs Manual | **-21.664** | **3.394e-14*** | 1.397 | 7.965260 | **4.805** | **4.284e-03*** |
| | | Baseline vs BMI | **-8.959** | **5.759e-07*** | **11.111** | **1.337e-08*** | **6.093** | **2.153e-04*** |
| | | Manual vs BMI | **9.891** | **1.056e-07*** | **7.638** | **7.718e-06*** | 0.118 | 40.816879 |
| | Beta (13-35Hz) | Baseline vs Manual | **67.642** | **2.078e-24*** | **31.919** | **1.254e-17*** | **8.690** | **9.590e-07*** |
| | | Baseline vs BMI | **56.723** | **8.219e-23*** | **33.253** | **5.389e-18*** | **11.422** | **8.093e-09*** |
| | | Manual vs BMI | 3.506 | 0.094663 | 1.770 | 4.105440 | -2.659 | 0.660268 |
| | Gamma (35-75Hz) | Baseline vs Manual | **33.551** | **4.485e-18*** | **10.681** | **2.715e-08*** | 3.121 | 0.232506 |
| | | Baseline vs BMI | **36.025** | **1.031e-18*** | 0.446 | 29.695306 | **-7.973** | **3.913e-06*** |
| | | Manual vs BMI | -3.667 | 0.064683 | **-5.369** | **1.133e-03*** | **-8.055** | **3.319e-06*** |
| | Highgamma (75-150Hz) | Baseline vs Manual | 1.343 | 8.718462 | **5.606** | **6.544e-04*** | **5.536** | **7.696e-04*** |
| | | Baseline vs BMI | -0.696 | 22.229325 | **-5.300** | **1.334e-03*** | -0.844 | 18.367808 |
| | | Manual vs BMI | -2.628 | 0.707687 | **-6.402** | **1.080e-04*** | **-6.064** | **2.304e-04*** |

**Supplementary Table 1. Relevant statistics for comparing average power at the go cue between each task type for each frequency band.** Paired t-test t-statistic and Bonferonni corrected p for comparison between average power normalized to total power during baseline vs manual control vs BMI control at the go cue for each monkey and frequency band. Significant differences are signified by bolded lettering and a (*).

**Target:** Comparing Average Power Between Task Types

| | | | M1 | | DLPFC | | Cd | |
|---|---|---|---|---|---|---|---|---|
| **Subject** | **Frequency Band** | **Comparison** | **t-statistic** | **Bonferroni corrected p** | **t-statistic** | **Bonferroni corrected p** | **t-statistic** | **Bonferroni corrected p** |
| Monkey H | Theta (4-8Hz) | Baseline vs Manual | 0.858 | 18.352398 | **9.584** | **2.546e-05*** | **-4.493** | **0.033084*** |
| | | Baseline vs BMI | -2.806 | 0.714292 | 3.336 | 0.266714 | -3.038 | 0.464430 |
| | | Manual vs BMI | **-8.563** | **8.379e-05*** | **-7.204** | **4.862e-04*** | 0.914 | 17.037005 |
| | Alpha (8-13Hz) | Baseline vs Manual | **8.470** | **9.383e-05*** | **6.627** | **1.099e-03*** | **6.193** | **2.087e-03*** |
| | | Baseline vs BMI | 3.812 | 0.111402 | 2.666 | 0.925721 | **8.321** | **1.129e-04*** |
| | | Manual vs BMI | **-14.092** | **3.563e-07*** | **-6.613** | **1.120e-03*** | 2.743 | 0.801803 |
| | Beta (13-35Hz) | Baseline vs Manual | **14.504** | **2.567e-07*** | **-4.276** | **0.048485*** | **7.408** | **3.683e-04*** |
| | | Baseline vs BMI | **11.521** | **3.421e-06*** | 0.257 | 36.069838 | **5.897** | **3.277e-03*** |
| | | Manual vs BMI | **-4.999** | **0.013939*** | **5.744** | **4.162e-03*** | -1.197 | 11.449104 |
| | Gamma (35-75Hz) | Baseline vs Manual | **-16.978** | **4.209e-08*** | **-9.743** | **2.133e-05*** | **-4.371** | **0.041011*** |
| | | Baseline vs BMI | **-13.338** | **6.645e-07*** | **-15.354** | **1.339e-07*** | **-12.653** | **1.203e-06*** |
| | | Manual vs BMI | **11.778** | **2.679e-06*** | **-7.277** | **4.403e-04*** | **-11.568** | **3.271e-06*** |
| | Highgamma (75-150Hz) | Baseline vs Manual | **-37.270** | **4.023e-12*** | -3.059 | 0.446434 | **-20.883** | **3.789e-09*** |
| | | Baseline vs BMI | **-25.543** | **3.551e-10*** | **-6.223** | **1.993e-03*** | **-18.878** | **1.230e-08*** |
| | | Manual vs BMI | **11.805** | **2.611e-06*** | -1.006 | 15.038327 | -2.252 | 1.973270 |
| Monkey Y | Theta (4-8Hz) | Baseline vs Manual | **-12.764** | **1.040e-09*** | **-4.988** | **2.772e-03*** | **-5.407** | **1.038e-03*** |
| | | Baseline vs BMI | **-23.752** | **5.285e-15*** | **-6.227** | **1.595e-04*** | **-7.282** | **1.617e-05*** |
| | | Manual vs BMI | -3.223 | 0.183528 | 3.039 | 0.280638 | 1.291 | 9.481099 |
| | Alpha (8-13Hz) | Baseline vs Manual | -2.979 | 0.322256 | 0.102 | 41.401543 | **5.198** | **1.693e-03*** |
| | | Baseline vs BMI | **-10.931** | **1.793e-08*** | -0.671 | 22.940993 | **5.696** | **5.317e-04*** |
| | | Manual vs BMI | **-7.863** | **4.884e-06*** | -0.852 | 18.180915 | -1.953 | 2.890856 |
| | Beta (13-35Hz) | Baseline vs Manual | **37.240** | **5.193e-19*** | **7.426** | **1.198e-05*** | **5.799** | **4.204e-04*** |
| | | Baseline vs BMI | **31.569** | **1.573e-17*** | **4.780** | **4.539e-03*** | **6.829** | **4.245e-05*** |
| | | Manual vs BMI | **-4.556** | **7.734e-03*** | **-6.934** | **3.387e-05*** | -2.866 | 0.416193 |
| | Gamma (35-75Hz) | Baseline vs Manual | 0.453 | 29.484801 | 2.972 | 0.327311 | 1.600 | 5.604214 |
| | | Baseline vs BMI | -2.029 | 2.489650 | **-10.386** | **4.470e-08*** | -3.073 | 0.259445 |
| | | Manual vs BMI | -1.908 | 3.154715 | **-11.266** | **1.039e-08*** | -2.883 | 0.400557 |
| | Highgamma (75-150Hz) | Baseline vs Manual | **-11.982** | **3.367e-09*** | **4.510** | **8.647e-03*** | **5.064** | **2.321e-03*** |
| | | Baseline vs BMI | **-15.470** | **2.667e-11*** | -3.389 | 0.124508 | **6.932** | **3.404e-05*** |
| | | Manual vs BMI | **3.949** | **0.033054*** | -8.501 | 1.380e-06 | -1.918 | 3.095394 |

**Supplementary Table 2. Relevant statistics for comparing average power at target acquisition between each task type for each frequency band.** Paired t-test t-statistic and Bonferonni corrected p for comparison between average power normalized to total power during baseline vs manual control vs BMI control at target acquisition for each monkey and frequency band. Significant differences are signified by bolded lettering and a (*).

**a**

Go Cue: Normalized Granger Causality vs 0

| Control Type | Subject | Interaction Direction | t-statistic | Bonferonni corrected p |
|---|---|---|---|---|
| BMI | Monkey H | M1→DLPFC | 0.183 | 10.294381 |
| | | **DLPFC→M1** | **6.779** | **2.353e-04*** |
| | | M1→Cd | -3.516 | 0.051088 |
| | | **Cd→M1** | **6.186** | **5.625e-04*** |
| | | DLPFC→Cd | -3.110 | 0.108243 |
| | | **Cd→DLPFC** | **6.763** | **2.409e-04*** |
| | Monkey Y | M1→DLPFC | -0.347 | 8.782855 |
| | | **DLPFC→M1** | **10.463** | **1.046e-08*** |
| | | **M1→Cd** | **-9.023** | **1.362e-07*** |
| | | **Cd→M1** | **14.460** | **2.612e-11*** |
| | | **DLPFC→Cd** | **-6.417** | **2.788e-05*** |
| | | Cd→DLPFC | 2.281 | 0.397076 |
| Manual | Monkey H | M1→DLPFC | 0.514 | 7.398460 |
| | | **DLPFC→M1** | **7.859** | **5.412e-05*** |
| | | M1→Cd | 0.439 | 8.018282 |
| | | **Cd→M1** | **4.229** | **0.014034*** |
| | | **DLPFC→Cd** | **5.527** | **1.565e-03*** |
| | | **Cd→DLPFC** | **-4.282** | **0.012782*** |
| | Monkey Y | **M1→DLPFC** | **-8.858** | **1.857e-07*** |
| | | **DLPFC→M1** | **12.398** | **4.774e-10*** |
| | | **M1→Cd** | **-7.839** | **1.367e-06*** |
| | | **Cd→M1** | **13.124** | **1.648e-10*** |
| | | **DLPFC→Cd** | **-3.873** | **0.010553*** |
| | | Cd→DLPFC | -1.447 | 1.951595 |

**b**

Target: Normalized Granger Causality vs 0

| Control Type | Subject | Interaction Direction | t-statistic | Bonferonni corrected p |
|---|---|---|---|---|
| BMI | Monkey H | M1→DLPFC | 0.180 | 10.321333 |
| | | **DLPFC→M1** | **5.573** | **1.454e-03*** |
| | | M1→Cd | -1.672 | 1.444127 |
| | | **Cd→M1** | **5.377** | **1.993e-03*** |
| | | DLPFC→Cd | -2.420 | 0.388041 |
| | | **Cd→DLPFC** | **5.418** | **1.865e-03*** |
| | Monkey Y | M1→DLPFC | 2.045 | 0.643538 |
| | | **DLPFC→M1** | **14.307** | **3.201e-11*** |
| | | **M1→Cd** | **-3.314** | **0.039562*** |
| | | **Cd→M1** | **11.708** | **1.374e-09*** |
| | | **DLPFC→Cd** | **-4.832** | **1.070e-03*** |
| | | **Cd→DLPFC** | **-8.625** | **2.898e-07*** |
| Manual | Monkey H | M1→DLPFC | 0.912 | 4.559055 |
| | | **DLPFC→M1** | **6.765** | **2.402e-04*** |
| | | M1→Cd | 2.985 | 0.136545 |
| | | **Cd→M1** | **3.734** | **0.034255*** |
| | | **DLPFC→Cd** | **5.556** | **1.495e-03*** |
| | | **Cd→DLPFC** | **-4.828** | **4.958e-03*** |
| | Monkey Y | M1→DLPFC | -0.514 | 7.349721 |
| | | **DLPFC→M1** | **9.897** | **2.785e-08*** |
| | | **M1→Cd** | **-2.701** | **0.160473*** |
| | | **Cd→M1** | **9.344** | **7.519e-08*** |
| | | **DLPFC→Cd** | **-4.613** | **1.801e-03*** |
| | | **Cd→DLPFC** | **-3.909** | **9.690e-03*** |

**Supplementary Table 3. Relevant statistics for comparing normalized granger causality to 0 for both BMI and manual control** (a) 1-sample t-test t-statistic and Bonferonni corrected p for comparison of normalized granger causality during BMI and manual control to 0 at go cue for each monkey and interaction direction. Significant differences are signified by bolded lettering and a (*). (b) Same as (a) but for target acquisition, rather than go cue.

**a**

Go Cue: BMI vs Manual Normalized Granger Causality

| Subject | Interaction Direction | t-statistic | Bonferonni corrected p |
|---|---|---|---|
| Monkey H | M1→DLPFC | -0.235 | 9.818311 |
| | DLPFC→M1 | 0.093 | 11.127620 |
| | **M1→Cd** | **-5.681** | **1.226e-03*** |
| | **Cd→M1** | **6.893** | **2.002e-04*** |
| | **DLPFC→Cd** | **-9.882** | **4.882e-06*** |
| | **Cd→DLPFC** | **10.185** | **3.525e-06*** |
| Monkey Y | **M1→DLPFC** | **8.213** | **6.474e-07*** |
| | DLPFC→M1 | -1.463 | 1.898857 |
| | M1→Cd | -0.799 | 5.197626 |
| | Cd→M1 | 2.303 | 0.379286 |
| | **DLPFC→Cd** | **-4.668** | **1.579e-03*** |
| | Cd→DLPFC | 2.813 | 0.125110 |

**b**

Target: BMI vs Manual Normalized Granger Causality

| Subject | Interaction Direction | t-statistic | Bonferonni corrected p |
|---|---|---|---|
| Monkey H | M1→DLPFC | -1.187 | 3.098871 |
| | DLPFC→M1 | 0.100 | 11.062261 |
| | **M1→Cd** | **-6.623** | **2.947e-04*** |
| | **Cd→M1** | **5.785** | **1.042e-03*** |
| | **DLPFC→Cd** | **-9.106** | **1.171e-05*** |
| | **Cd→DLPFC** | **10.515** | **2.492e-06*** |
| Monkey Y | M1→DLPFC | 2.743 | 0.146179 |
| | **DLPFC→M1** | **5.388** | **2.896e-04*** |
| | M1→Cd | -1.366 | 2.234999 |
| | Cd→M1 | 1.531 | 1.689782 |
| | DLPFC→Cd | -1.275 | 2.596613 |
| | **Cd→DLPFC** | **-3.646** | **0.018125*** |

**Supplementary Table 4. Relevant statistics for comparing between normalized granger causality during BMI and manual control** (a) Paired t-test t-statistic and Bonferonni corrected p for comparison between normalized granger causality during BMI vs manual control at go cue for each monkey and interaction direction. Significant differences are signified by bolded lettering and a (*). (b) Same as (a) but for target acquisition, rather than go cue.

**a**

Go Cue: Net Normalized Granger Causality vs 0

| Control Type | Subject | Interaction Direction | Net Normalized GC | SEM Across Days | t-statistic | Bonferonni corrected p |
|---|---|---|---|---|---|---|
| BMI | Monkey H | **DLPFC→M1** | **0.737** | **0.137** | **5.392** | **9.734e-04*** |
| | | **Cd→M1** | **1.294** | **0.226** | **5.715** | **5.810e-04*** |
| | | **Cd→DLPFC** | **0.383** | **0.082** | **4.672** | **3.236e-03*** |
| | Monkey Y | **DLPFC→M1** | **0.088** | **0.017** | **5.304** | **1.762e-04*** |
| | | **Cd→M1** | **0.232** | **0.019** | **12.167** | **3.386e-10*** |
| | | **Cd→DLPFC** | **0.157** | **0.019** | **8.456** | **2.008e-07*** |
| Manual | Monkey H | **DLPFC→M1** | **0.728** | **0.111** | **6.556** | **1.624e-04*** |
| | | Cd→M1 | 0.535 | 0.192 | 2.791 | 0.097935 |
| | | **Cd→DLPFC** | **-0.605** | **0.113** | **-5.366** | **1.015e-03*** |
| | Monkey Y | **DLPFC→M1** | **0.163** | **0.013** | **12.205** | **3.196e-10*** |
| | | **Cd→M1** | **0.210** | **0.019** | **11.314** | **1.283e-09*** |
| | | Cd→DLPFC | 0.033 | 0.015 | 2.228 | 0.221623 |

**b**

Target: Net Normalized Granger Causality vs 0

| Control Type | Subject | Interaction Direction | Net Normalized GC | SEM Across Days | t-statistic | Bonferonni corrected p |
|---|---|---|---|---|---|---|
| BMI | Monkey H | **DLPFC→M1** | **0.562** | **0.126** | **4.457** | **4.701e-03*** |
| | | **Cd→M1** | **0.987** | **0.221** | **4.473** | **4.573e-03*** |
| | | **Cd→DLPFC** | **0.297** | **0.087** | **3.402** | **0.031479*** |
| | Monkey Y | **DLPFC→M1** | **0.094** | **0.015** | **6.177** | **2.379e-05*** |
| | | **Cd→M1** | **0.147** | **0.022** | **6.841** | **5.521e-06*** |
| | | Cd→DLPFC | **-0.032** | **0.016** | -2.044 | 0.322138 |
| Manual | Monkey H | **DLPFC→M1** | **0.539** | **0.104** | **5.179** | **1.379e-03*** |
| | | Cd→M1 | 0.233 | 0.194 | 1.200 | 1.520397 |
| | | **Cd→DLPFC** | **-0.633** | **0.115** | **-5.496** | **8.231e-04*** |
| | Monkey Y | **DLPFC→M1** | **0.083** | **0.015** | **5.465** | **1.210e-04*** |
| | | **Cd→M1** | **0.121** | **0.021** | **5.739** | **6.424e-05*** |
| | | Cd→DLPFC | 0.006 | 0.014 | 0.435 | 4.008821 |

**Supplementary Table 5. Relevant statistics for comparing net normalized granger causality to 0 for both BMI and manual control** (a) Net normalized Granger causality values and standard error mean (SEM) across days at go cue as well as t-statistic and Bonferonni corrected p from 1-sample t-tests comparing net normalized granger causality during BMI and manual control to 0 at go cue for each monkey and interaction direction. Significant differences are signified by bolded lettering and a (*). (b) Same as (a) but for target acquisition, rather than go cue.
